# Decomposition disentangled: a test of the multiple mechanisms by which nitrogen enrichment alters litter decomposition

**DOI:** 10.1101/671545

**Authors:** Noémie A. Pichon, Seraina Cappelli, Santiago Soliveres, Norbert Hölzel, Valentin H. Klaus, Till Kleinebecker, Eric Allan

**Author notes:** Corresponding author: Noémie A. Pichon.

## Abstract

1. Nitrogen (N) enrichment has direct effects on ecosystem functioning by altering soil abiotic conditions and indirect effects by reducing plant diversity and shifting plant functional composition from dominance by slow to fast growing species. Litter decomposition is a key ecosystem function and is affected by N enrichment either by a change in litter quality (the recalcitrance of the plant material) or through a change in soil quality (the abiotic and biotic components of the soil that affect decomposition). The relative importance of soil and litter quality and how the direct and effects of N alter them remains poorly known.
2. We designed a large grassland field experiment manipulating N enrichment, plant species richness and functional composition in a full factorial design. We used three complementary litter bag experiments and a novel structural equation modelling approach to quantify the relative effects of the treatments on litter and soil quality and their importance for total decomposition.
3. Our results indicate that total decomposition was mostly driven by changes in litter quality rather than soil quality. Litter quality was affected by the nutrient contents (N and calcium) and structural components of the litter (leaf dry matter content, fibres). N enrichment increased litter decomposition mostly indirectly through a shift in functional composition toward faster growing plant species producing higher quality litter. N enrichment also had effects on soil, by directly and indirectly affected vegetation cover, but this had relatively few consequences for the total decomposition rate.
4. *Synthesis*. Our approach provides a mechanistic tool to test the drivers of litter decomposition across different ecosystems. Our results show that functional composition is more important than richness or soil quality in determining litter decomposition and that N enrichment effects mainly occur via above- rather than belowground processes. This highlights the importance of considering shifts in plant species composition when assessing the effects of N enrichment on decomposition.

## Introduction

Soil nitrogen enrichment is one of the major global changes ecosystems are currently facing (Galloway et al., 2008). Nitrogen (N) enrichment alters ecosystem functioning directly and through several indirect mechanisms. It directly alters functions related to nutrient stocks and fluxes by changing soil abiotic conditions, stoichiometry and pH (Sardans, Rivas-Ubach, & Peñuelas, 2012; Laliberté & Tylianakis, 2012). In addition N enrichment indirectly affects ecosystem functioning by altering biotic community properties such as plant diversity and composition. N enrichment typically reduces the number of plant species able to coexist (Suding et al., 2005) and this loss of diversity could affect ecosystem functioning as much as N addition per se. (Hooper et al., 2012; Tilman, Reich, & Isbell, 2012). However, plant community change, following N enrichment, does not only involve a loss of species it also involves compositional turnover and in particular a shift towards faster growing plant species (Isbell et al., 2013; Lavorel & Grigulis, 2012; de Vries et al., 2012). This shift is indicated by an increase in mean values of trait linked to the leaf economics spectrum, such as specific leaf area and leaf N content, (Wright et al., 2004) and this shift is a key driver of ecosystem functioning (Lavorel & Grigulis, 2012). However, we still have little mechanistic insight into the relative importance of these direct (abiotic) and indirect (plant richness and composition) effects of N enrichment on ecosystem functioning. Observational studies have separated direct effects of N from indirect effects mediated through species richness (Isbell et al., 2013) and/or functional composition (Allan et al., 2015). However, observational studies struggle to separate effects of correlated drivers, such as diversity loss and compositional turnover. Experimental approaches are therefore needed to separate these effects and to fully understand and predict the mechanisms by which N addition affects ecosystem functioning.

The decomposition of plant litter is a key ecosystem function that influences rates of soil biogeochemical cycling and which is strongly affected by N deposition (Finn et al., 2015; Knorr, Frey, & Curtis, 2005; Hobbie et al., 2012). Depending on the ecosystem, the enrichment level and duration, N can have either positive or negative effects on decomposition (Bardgett & Wardle, 2012; Knorr et al., 2005; Hobbie et al., 2012; Riggs, Hobbie, Bach, Hofmockel, & Kazanski, 2015) and to understand this variation we need to better understand the mechanisms behind them. Plant litter decomposition is determined by multiple mechanisms: it depends principally on the physical and chemical properties of the litter and on soil biotic and abiotic conditions (Cebrian, 1999; Handa et al., 2014; Cornwell et al., 2008). To distinguish these two main drivers of litter decomposition, we will refer to “litter quality”, as the physical and chemical properties of litter that affect its decomposition and to “soil quality”, as the soil biotic and abiotic factors which determine decomposition rates. Both soil and litter quality are key determinants of litter decomposition but their relative importance, especially following N enrichment, is not well known (but see Cleveland et al., 2014; García-Palacios, Prieto, Ourcival, & Hättenschwiler, 2016b; Maaroufi, Nordin, Palmqvist, & Gundale, 2017). N enrichment could influence decomposition by directly or indirectly changing both soil quality (i.e. by altering soil properties and fauna), and litter quality. To understand the impacts of N enrichment on decomposition we need experimental and analytical approaches that can separate these different, cascading mechanisms.

N enrichment is likely to directly and indirectly alter litter quality and therefore decomposition rates. Litter quality is largely determined by its chemical properties (nutrient contents and the presence of defence compounds) and by physical factors such as leaf dry matter and fibre contents (Garnier et al., 2004; Cornwell et al., 2008). With higher soil N availability, plants will produce more rapidly degradable tissues with higher N contents and fewer fibres. In addition to N, macronutrients like Ca and Mg may also influence litter decomposability (García-Palacios, McKie, Handa, Frainer, & Hättenschwiler, 2016a) and their availability could also be altered by N addition (Aber et al., 1998). Indirect effects of N are also likely to be important: a shift to fast growing plant communities further enhances litter quality because fast growing plants have generally higher leaf N and lower fibre contents. Fast growing plants also invest less in defences against herbivores and pathogens (Blumenthal, Mitchell, Pysek, & Jarosík, 2009) and have fewer chemicals such as tannins that reduce decomposition. However, other indirect effects of N may reduce decomposition. A reduction in species and functional diversity could reduce decomposability (Handa et al., 2014). Although some aspects of litter quality are well characterised, we lack a comprehensive picture of how N enrichment alters these different aspects simultaneously.

Enriching soils with N is likely to alter their quality for litter decomposition both directly and indirectly. The abundance and composition of the soil macro, meso and microfauna are key determinants of soil quality (Milcu & Manning, 2011) as macrofauna fragment large litter pieces, which accelerates decomposition by smaller organisms (Milcu, Partsch, Scherber, Weisser, & Scheu, 2008). N enrichment could increase soil quality if it causes a shift towards bacterial dominated communities (from fungal dominated ones), either through direct effects of N or through changes in plant functional composition, which is likely to lead to increased decomposition rates (Fierer, Strickland, Liptzin, Bradford, & Cleveland, 2009; Bardgett & McAlister, 1999; Bardgett & Wardle, 2012; de Vries, Hoffland, van Eekeren, Brussaard, & Bloem, 2006). However, N enrichment might indirectly reduce soil quality if a loss of plant diversity loss results in a loss of soil organism diversity (Milcu et al., 2013). In addition, N addition will directly increase plant biomass (in N limited systems), but might indirectly reduce it by reducing diversity (van der Plas, 2019; Isbell et al., 2013), and a change in biomass will alter microclimatic conditions such as soil temperature and moisture, which are important drivers of decomposition (Hättenschwiler, Tiunov, & Scheu, 2005; Blankinship, Niklaus, & Hungate, 2011). The various direct and indirect effects of N enrichment are therefore likely to have complex and potentially opposing effects on soil quality and therefore on litter decomposition rates.

In this study, we tested the effects of N enrichment on litter decomposition and disentangled its direct effects on soil and litter quality from its indirect effects mediated by plant richness and functional composition. We created experimental plant communities to realise a full factorial cross of plant functional composition, plant species richness and N enrichment. Plant functional composition was manipulated by creating a gradient in community mean specific leaf area and leaf N as these traits are key indicators of resource economics and plant growth strategy. Three complementary litter bag experiments were used to test direct and indirect effects of N addition on litter quality, on soil quality and on both combined. We also looked at the effect of macro and mesofauna on decomposition by using different mesh sized litter bags. This framework enabled us to test the following questions:

What is the relative importance of direct effects of N enrichment on decomposition relative to indirect effects mediated through changes in the plant community (species richness and functional composition)?

Is decomposition determined more by changes in litter quality or soil quality?

How important are meso and macro fauna in determining decomposition and how does their relative importance change with N enrichment?

## Material and methods

### The PaNDiv Experiment

The PaNDiv Experiment is located in Münchenbuchsee near the city of Bern (Switzerland, 47°03’N, 7°46’E, 564 m.a.s.l.). It has a mean annual temperature of 9.2 ± 0.61°C and mean annual precipitation of 1051.78 ± 168.42 mm y^−1^ (mean over the last 30 years, data from the Federal Office of Meteorology and Climatology MeteoSwiss). The soil is characterized as 0.7 to 1m deep brown earth (Cambisol), according to the Geoportal of the Canton Bern (http://www.geo.apps.be.ch). We measured total soil N and carbon (C) concentrations and pH in the top 20 cm of soil at the start of the experiment and found concentrations of 2.3-4.2% C, 0.26-0.43% N and a pH of 7.4. The field site had been extensively managed (without fertilization) for at least 10 years before the start of the experiment and had been used for fodder production and grazing. The vegetation was cleared and the area ploughed before the experimental plots were established.

The species sown were selected from a pool of 20 species commonly found in both extensively and intensively managed Central European grasslands. We divided our 20 species into 10 fast and 10 slow growing species according to their Specific Leaf Area (SLA) and leaf N content, which are related to resource use strategy (see Figures S1 and S2) (Wright et al., 2004). The fast growing pool therefore corresponds to species found in N enriched sites, whereas the slow growing pool comprises species found in less productive sites. We excluded legumes from the species pool as few legume species will grow well at high N levels and including legumes only in the slow growing pool would have caused an additional and large difference between the species pools. We realised several combinations of fast and slow growing species, so effects of changes in mean traits are independent of particular species effects.

In order to separate direct and indirect effects of N enrichment, we established a factorial cross of treatments representing the direct (N enrichment) and indirect effects (plant diversity loss and change in functional composition) on 2×2m plots. Fertilised plots received N in the form of urea twice a year in April and late June (beginning of the growing season and following the first cut, see below), for an annual addition of 100 kg N ha^−1^y^−1^, which corresponds to intermediately intensive grassland management (Blüthgen et al., 2012). To manipulate diversity, we established plots with 1, 4, 8 or all 20 species. To manipulate functional composition and diversity we established plots with only fast growing, only slow growing or a mix of fast and slow growing species. This allowed us to realise a large gradient in community weighted mean trait values, which is crossed with functional diversity, as mixed plots have higher diversity than single strategy plots. Functional composition, functional diversity and species richness were all completely crossed at the 4 and 8 species levels (monocultures and 20 species plots could only contain one functional composition). We sowed all plants in monoculture and we established four replicates of the 20 species together. At the four and eight species levels we randomly selected species compositions: we selected 10 species compositions for each combination of richness (4 and 8), times functional composition (fast, slow mixed). This meant we had a total of 20 monocultures, 30 four species compositions, 30 eight species compositions and four replicates of 20 species composition. We constrained the random selection to ensure that all polycultures contained both grasses and herbs. The 84 different species compositions were grown once in control conditions and once with N addition. In addition to the N treatment, we also applied a fungicide treatment and a fungicide x N treatment, resulting in 336 plots in total. However, for logistical reasons the litter bag experiment was only conducted on the 168 control (no fungicide) plots (see Table S1). The whole field was divided into four blocks. Each block contained all 84 compositions but the particular N x fungicide treatment was randomly allocated per block. A regularly mown 1m path sown with a grass seed mixture consisting of *Lolium perenne* and *Poa pratensis* (UFA-Regeneration Highspeed) separated the plots.

All species within a plot were sown at equal density in October 2015, with proportions corrected by species specific germination rates, to obtain a total density as close as possible to 1000 seedlings m^−2^. The seeds were obtained from commercial suppliers (UFA Samen, Switzerland, and Rieger-Hofmann, Germany). Some species were resown once in spring 2016 because of poor establishment (*Heracleum sphondylium*, *Anthriscus sylvestris*, *Daucus carota*, *Salvia pratensis*, *Prunella grandiflora*, *Plantago media*), because they were mixed with other seeds to begin with (*Helictotrichon pubescens*, *Bromus erectus*) or because their seedlings froze in autumn or spring (*Holcus lanatus*, *Dactylis glomerata*, *Anthoxanthum odoratum*). No resowing was done after spring 2016. In order to maintain the diversity levels, the plots were weeded three times a year in April, July and September. This regime was highly successful and most plots contained very low weed covers in the following season (Figure S3). The whole experiment was mown twice a year in mid-June and mid-August which corresponds to intermediate to extensive grassland management.

### Measuring decomposition of litter bags

We conducted three complementary litter bag experiments simultaneously to test the mechanisms by which our treatments affected decomposition. The first set of bags tested the effect of our treatments on the soil quality. We filled those bags with rapeseed straw (*Brassica napus*) as a standard material and placed them on every plot. No Brassicaceae are present in the experiment and this litter should therefore be equally foreign for all plots. To test the effect of our treatments on litter quality (decomposability), we filled a second set of bags with biomass collected from each plot and let them decompose in a common garden, established in the grassland surrounding the experimental plots. We filled the third set of bags, called plot bags, with aboveground dry biomass from each plot and let them decompose on their own plot (i.e. the plot from which the biomass was collected) to test the combined effect of soil and litter quality on decomposition. By combining data from these three experiments, we can disentangle the relative importance of soil and litter quality in driving overall decomposition.

We sewed the litterbags using nylon fabric with a mesh size of 5 mm for the above part and 0.2 mm for the fabric in contact with the soil, to avoid loss of material during transport and manipulation (Bradford, Tordoff, Eggers, Jones, & Newington, 2002). To investigate the effects of different sized groups of detritivores on decomposition, we sewed two additional plot bags: a 2 mm mesh size to exclude the macrofauna, and a 0.2 mm mesh size to exclude meso and macrofauna (Milcu & Manning, 2011; Bardgett, 2005). By comparing decomposition rates in the bags with different mesh sizes we can estimate the effect of different aspects of the soil community on the overall decomposition rate.

The plant biomass used to fill the common garden and plot bags was collected on the field before the mowing in June 2017 (with some very unproductive plots sampled again in August in order to have enough material). Green litter differs in its composition from senescent litter due to nutrient resorption (Aerts, 1996), and therefore decomposes at a different rate (Sanaullah, Chabbi, Lemaire, Charrier, & Rumpel, 2010). We were, however, more interested in the difference in decomposition among plant communities rather than in measuring the absolute decomposition rate. In addition, green litter decomposition is an important process in grasslands which are managed by cutting and many similar decomposition experiments have therefore also used green litter (Sanaullah et al., 2010; Vogel, Eisenhauer, Weigelt, & Scherer-Lorenzen, 2013). The biomass was dried at 65°C for 48h, chopped, homogenized and split into equal parts (Biomass splitter, RT 6.5–RT 7; Retsch, Haan, Germany). We filled each bag with a maximum of 20g dry material and weighed the litterbags again after closing. Because some experimental communities produced only a small amount of biomass, we could not include 20g in all bags and the initial biomass varied from 5 to 20g. The bags decomposed on top of the soil for 2.5 months between September and December 2017. We then collected the bags, cleaned them of debris and soil, dried them and weighed them again. We measured decomposition rate as the percentage biomass lost between September and December, to correct for differences in initial weight. Initial bag weight was included as a covariate in our models but it never affected the percentage mass loss (see Table S3).

### Plant traits used to calculate functional composition

To produce a continuous measure of functional composition for all our plant communities we calculated community weighted means for Specific Leaf Area (SLA) and Leaf Dry Matter Content (LDMC). Although plots were designed to differ in SLA, we also created a large gradient in mean LDMC, which was only partially correlated with SLA. We measured SLA and LDMC in the control (unfertilised) monocultures and therefore did not include any plasticity in response to N addition, in order to ensure that the community weighted mean traits were as orthogonal to N addition as possible. We sampled one leaf from five individuals per species and followed the protocol of Garnier, Shipley, Roumet, and Laurent (2001) and measured the fresh weight and leaf area with a leaf area meter (LI-3000C, LI-COR Biosciences) after a minimum of 6h and a maximum of 2 days of rehydration in the dark. We dried the samples at 65°C for two days and measured their dry weight. To measure the abundances of the plant species, we visually estimated the percentage cover of our target and weed species on every plot before the biomass was cut. In total three people estimated cover but there was no systematic difference in the species relative covers estimated by the three recorders (data not shown). We calculated a Community Weighted Mean (CWM) trait measure for each plot by multiplying each species’ relative abundance (cover) by the mean trait value of the species in monoculture (CWM = ∑ p_i_ * x_i_; with p_i_ the relative abundance of the species i and x_i_ the trait value of i).

### Litter quality

Two key aspects of litter quality are nutrient and fibre contents. We measured the concentration of several nutrients and fibre fractions in the plant biomass. We analysed biomass samples of all plots from June and August 2017 using Near Infrared Reflectance Spectrometry (NIRS). A minimum of 5 g of biomass per plot (pooled sample, including all species present and their relative abundance) was ground with a cyclone mill to obtain a fine powder. The infrared spectrum of the powder was used to estimate the nutrient and fibre contents based on calibration models developed for aboveground grassland biomass by Kleinebecker, Klaus, and Hölzel (2011). We estimated acid detergent fiber (ADF: cellulose, lignin and silica), neutral detergent fiber (NDF: ADF + hemicellulose) and acid detergent lignin (ADL: crude lignin fraction) in the biomass, as well as concentrations of N, C, phosphorus (P), potassium (K), calcium (Ca) and magnesium (Mg).

We could not use all nutrients and fibre fractions separately in the analyses as some of them were highly correlated (e.g. Mg and Ca, see Figures S4 and S5). We decided to select a widely used set of variables that did not correlate strongly and which together account for structural components and nutritional quality of litter: biomass N, fibres (ADF) and Ca content (García-Palacios et al., 2016a; Smith & Bradford, 2003; Cornwell et al., 2008). We did not include ratios like C:N or lignin:N as they were closely correlated with other variables and did not add more information to the model.

In addition to our measures of functional composition (CWMs) and mean values of litter quality, we calculated a measure of litter quality diversity. For this we used the abundance weighted Mean Pairwise Distance metric (MPD) (de Bello, Carmona, Lepš, Szava-Kovats, & Pärtel, 2016). This measure quantified the distance between all species in a plot in their SLA, LDMC, biomass N, fibre and Ca values. In order to derive species specific values for biomass N, Ca and fibres, we used the values from the control monocultures as the species trait values, as for SLA and LDMC.

### Analyses

We first used linear mixed effect models to test the effect of our treatments on litter decomposition (percentage mass loss), for each bag individually and for all sets of bags combined. We ran two combined models: one with plot litter, standard litter and common garden litter combined and one with the three mesh sizes combined. We ran the models in R (package lme4, Bates, Mächler, Bolker, & Walker, 2015; R Core Team, 2018) and simplified full models by dropping terms that did not significantly improve the overall model fit, using likelihood-ratios. All models included block and species composition (84 levels) as random terms. Species composition distinguished the randomly assembled sets of species and was included to correct for the fact that replicated species composition are pseudoreplicates for testing the species richness effects. The combined model with all the bags also included plot as a random term (168 levels). We added fixed covariates for the month of biomass harvest (June or August) and the initial weight of biomass put in each bag. We did not transform the data since the errors were normally distributed and the variance homogenous.

The first type of models tested the effects of the treatments on each bag:

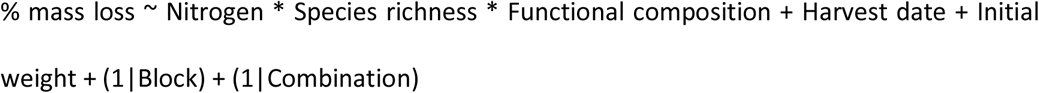

The second type of models tested for interactions between bag type (plot, standard, common garden litter; or the three mesh sizes) and the treatments:

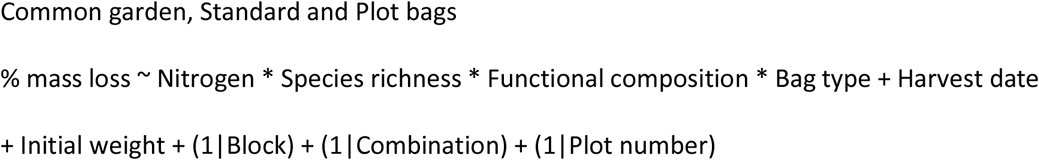

Although we used categorical measures of functional composition to design the experiment, we intended to create a gradient in CWM traits. We therefore replaced our three level functional composition variable by a continuous measure of community weighted mean SLA and LDMC, and functional diversity (MPD). For instance, in a single model:

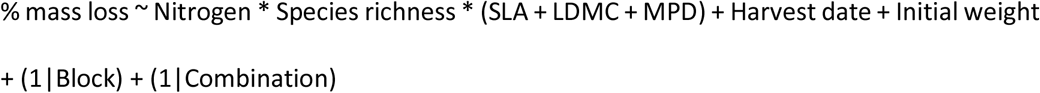

In a second step, we quantified the mechanisms by which our treatments affected decomposition using Structural Equation Modelling (SEM) (Grace, 2006). We included our three decomposition experiments (and the different mesh size treatments, see below) in the same model. By doing this we were able to test, not only the effect of our treatments on litter or soil mediated decomposition, but also the relative importance of litter and soil mediated decomposition for driving the final decomposition rate measured per plot. We used the mass loss in the “plot” litter bags (i.e. litter decomposing on its own plot) as a measure of the total plot decomposition rate. We then used the mass loss in the common garden litter bags as a measure of the litter mediated effects on decomposition, as these bags decompose on the same soil and only variation in litter quality will determine variation in mass loss between the bags. We used mass loss from the standard litter bags as our measure of soil mediated decomposition rates. In these bags the litter is always the same and therefore only variation in soil quality between plots will determine variation in decomposition. We fitted paths from common garden and standard litter mass loss to plot litter mass loss. The size of these two standardised path coefficients indicates the relative contribution of litter and soil quality to overall decomposition rates. In the SEM, plot litter mass loss is only affected by the mass loss measured in common garden and standard litter bags, to determine if we can explain all of the variation in overall decomposition rate based on our two measures of litter and soil quality.

We then tried to identify the traits and community properties that determined litter and soil quality. To do this we included our manipulated variables, N addition and plant species richness, as well as continuous measures of plant functional composition and litter quality, SLA, LDMC, biomass N, fibres and Ca, in the SEM. These measures could affect functional diversity (MPD) and microclimate. The microclimate measure we used in the analyses is the total plant cover on each plot. It correlates with biomass production and accounts for humidity and temperature variation among plots (Figure S6). To account for an effect of the soil fauna on decomposition, we included the log response ratio of the big mesh to the small mesh bag decomposition rate (see Figure S7). This variable “soil fauna effect” measures the relative effect of macro and mesofauna exclusion on decomposition and tests whether our treatments alter their effect.

We fitted SEMs using the lavaan package (Rosseel, 2012). This meant we could not include random effects, which could bias paths from species richness to other variables (which are not corrected for species composition). However, we also fitted models using piecewiseSEM (Lefcheck, 2016), in which we could include composition as a random effect, and this did not change the significance of any paths (see Table S2). Our initial model was rejected. We therefore included four residual covariance terms suggested by the lavaan modification indices. Including these covariances substantially improved model fit and led to a well supported model, however, it did not change the significance or substantially alter the strength of any paths. The first additions were negative covariances between biomass N and MPD and between LDMC and MPD. These are justified because monocultures (coded as zero MPD) had a greater range in biomass N and LDMC measures than polycultures, meaning some monocultures had much higher biomass N content than any of the polycultures. The two other covariances were between soil fauna effect and plot decomposition, and between litter quality and plot decomposition. These covariances are reasonable because we used the same litter in these different bags and a residual covariance is therefore likely. The residual covariance between plot and litter quality was fitted alongside a directed path and indicated the influence of unmeasured variables on both terms. The theoretical model and all detailed hypothesis are described in the Supplementary Information (Figure S7).

## Results

### Individual effects of N enrichment and plant community characteristics on litter and soil quality

Decomposition rates differed significantly among bag types. Litter decomposed faster in the common garden than on the experimental plots, and standard litter decomposed most slowly. Decomposition rates increased with mesh size (Fig.1a and Table S3).

N enrichment increased the litter decomposition rate in all bags consistently (significant main effect of N but no interaction between N and mesh size, Table S3 and Figure 1a). The effect was absent for standard litter bags when analysed alone but was significant when different bag types (common garden, standard, and plot big mesh size bags) were analysed together. There was no interaction between N and mesh size, meaning that N enrichment did not change the relative effect of large fauna, compared to small fauna, on decomposition.

**Figure 1.**
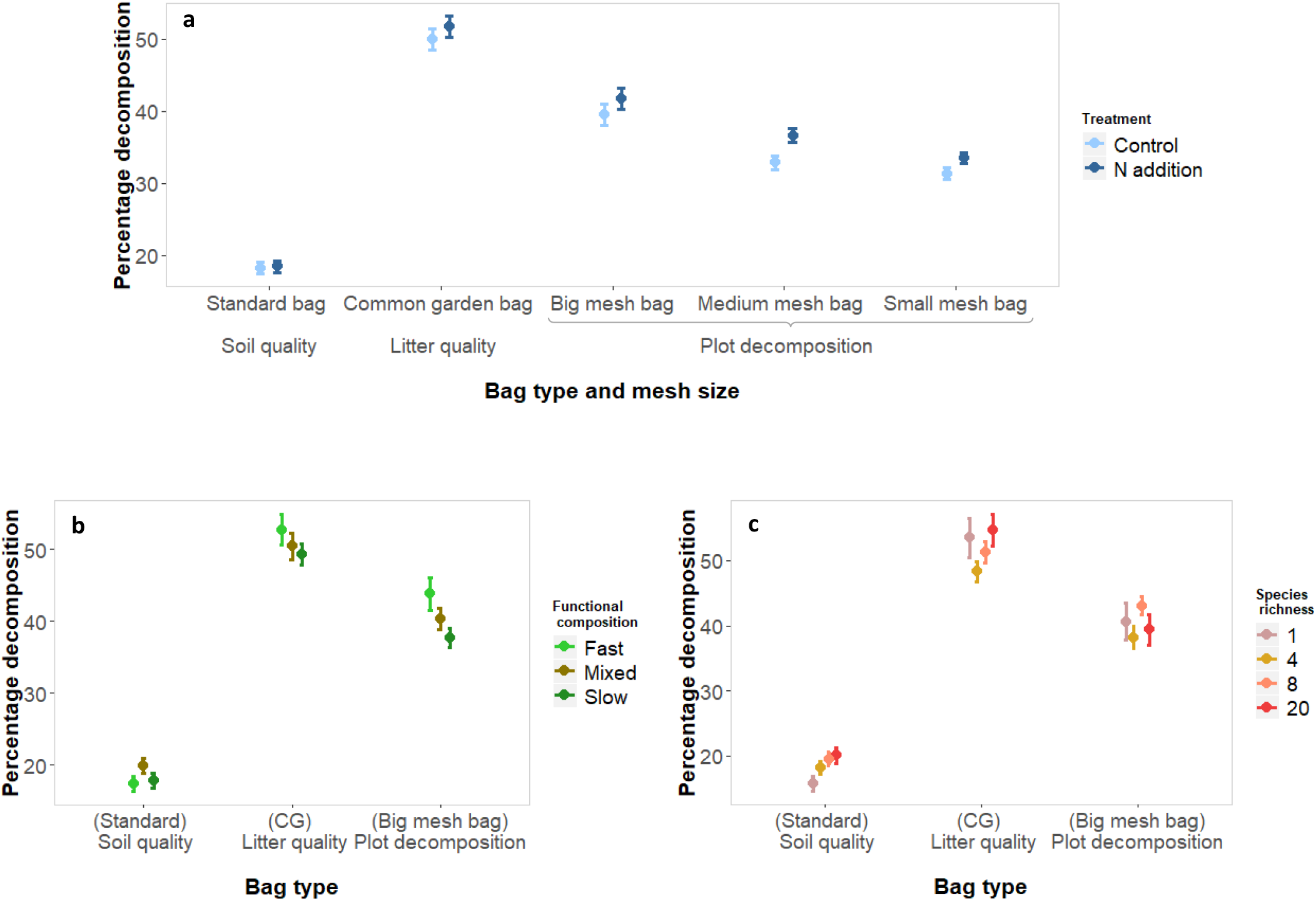
Effect of nitrogen addition (**a**), functional composition (**b**) and species richness (**c**) on litter decomposition depending on the litter bag type (standard, common garden and plot decomposition) and the mesh size (big, medium and small). Mean and standard error of the raw values (168 plots per bag).

Plant functional composition, expressed as a categorical variable (fast, mixed or slow growing species, Fig.1b), had a significant effect on the decomposition of common garden and plot litter. Litter from fast growing communities decomposed more rapidly than litter from mixed and slow communities. We observed the same pattern with continuous measures of functional composition, with a non-significant effect of SLA but a negative significant effect of LDMC on decomposition (Figure 2). LDMC therefore seemed to be a better predictor than SLA of the effect of growth strategy on decomposition. Comparing the bags with different mesh sizes, LDMC had a larger negative effect on decomposition in the big mesh litter bags than in the smaller mesh sizes, suggesting a larger effect of LDMC on the activity of the macrofauna than on the activity of the meso or microfauna (Fig 2b).

**Figure 2.**
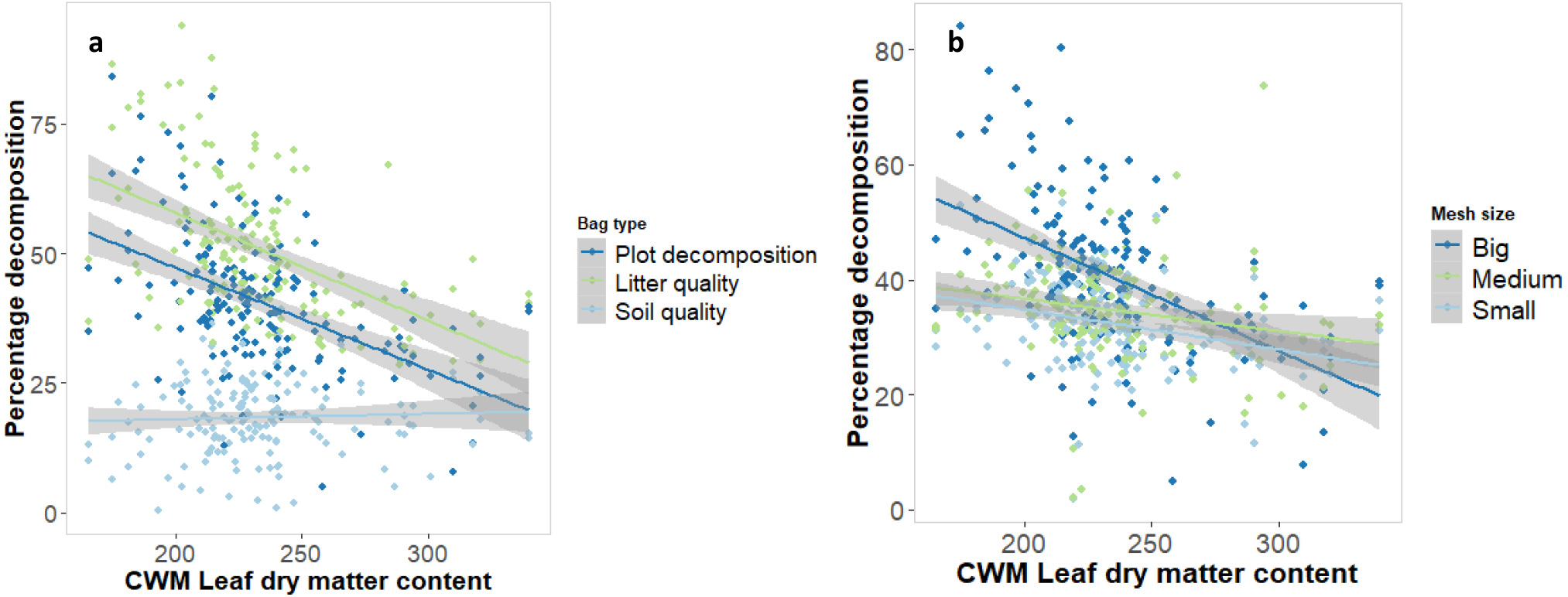
Effect of community weighted mean leaf dry matter content (mg g^−1^) on decomposition depending on the bag type (**a**) and on the mesh size (**b**). Mean and standard error of the raw values (168 plots per bag).

Plant species richness had a positive effect on the decomposition of standard litter bags, when analysed separately (Fig.1c and Table S3). The effect of functional diversity depended on the bag type, with a negative effect in plot and common garden bags and a positive effect on standard bags (Table S3). These results indicate that species richness and functional diversity of communities increased soil quality, whereas the functional composition of the community increased litter quality.

### Relative importance of litter and soil quality in driving overall decomposition

We used structural equation models to test the relative importance of our different treatments in affecting soil and litter quality and the relative importance of litter and soil in driving the overall decomposition rate. Litter and soil quality both had a positive effect on total plot decomposition, but litter quality was much more important (path coefficient of 0.96, Table S4) than soil quality (path coefficient of 0.20; see Figures 3 and 4a and b). Although soil macro and mesofauna increased decomposition overall, they did not contribute to variation in decomposition between plots, as there was no link between the log response ratio between decomposition in big and small mesh-sized bags and the overall decomposition rates.

**Figure 3.**
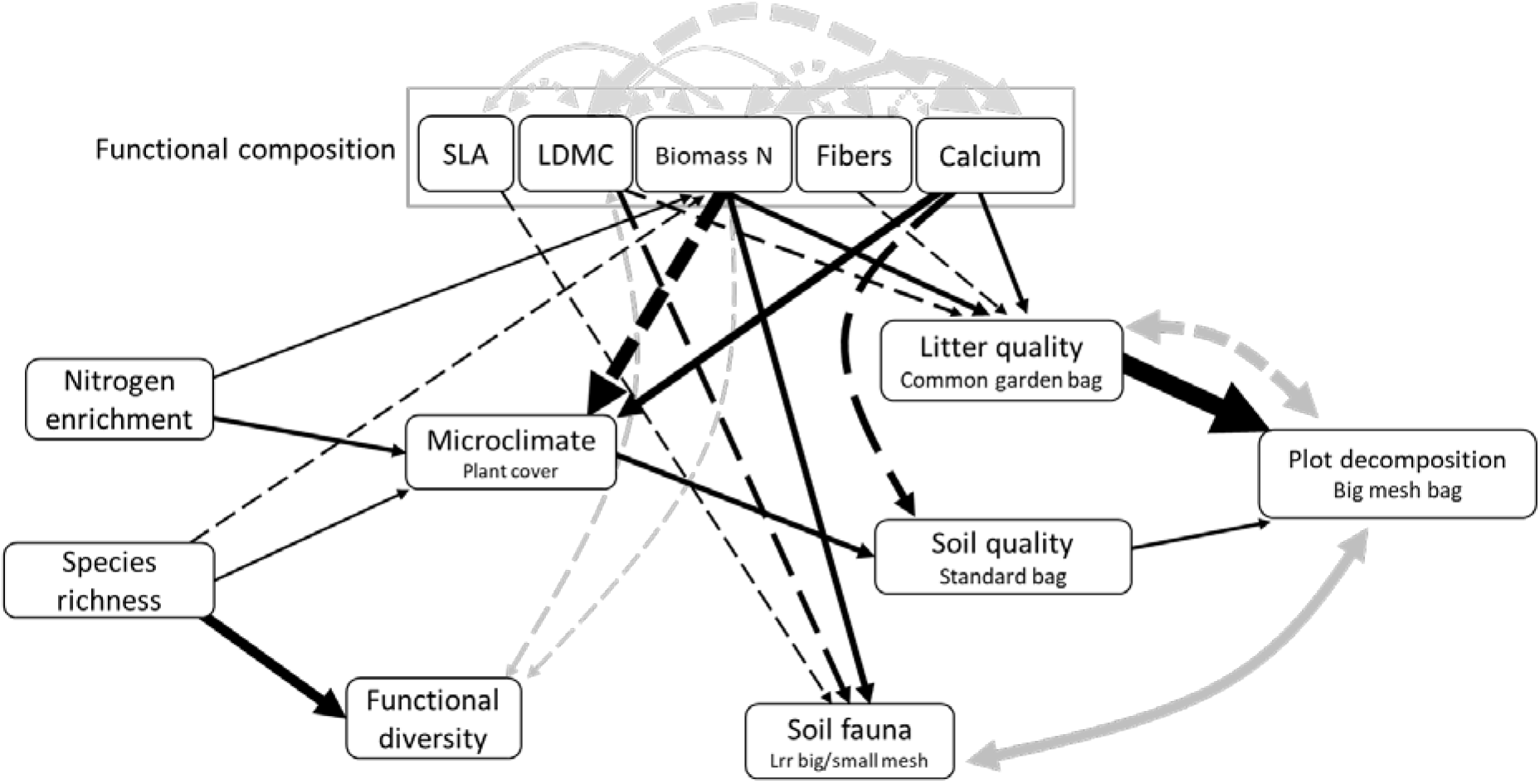
Final results of the structural equation model, showing effects of nitrogen enrichment, plant species richness and plant functional composition on decomposition. Dashed arrows show negative, full arrows positive path coefficients. The arrow size is proportional to the path coefficient. Double-headed grey arrows show covariances. Details of the output in Table S4. Model fit: Pvalue 0.423; Chisq 17.477; Df 17; RMSEA 0.013.

**Figure 4.**
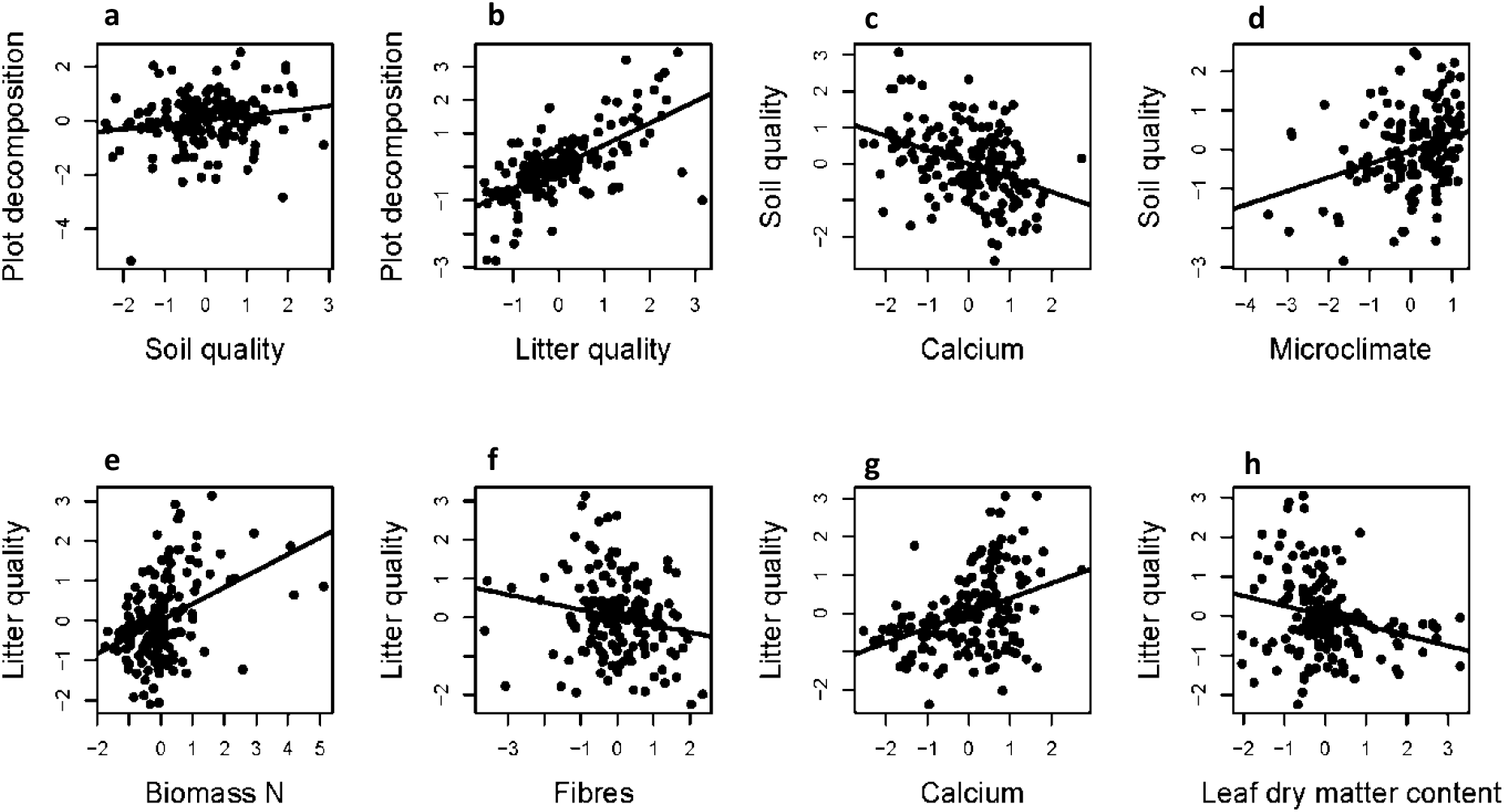
Partial plots visualizing SEM outputs from the Figure 3 of variables effects on overall decomposition (**a-b**), on soil quality (**c-d**) and on litter quality (**e-h**). X-axis units are standardized values, y-axis are standardized residuals of the target explanatory variable on the remaining explanatory variables.

Litter quality was mainly influenced by plant functional composition. Litter from communities with a high biomass N content, low LDMC and low fibre content, corresponding to our fast growing communities, decomposed faster than litter from slow growing communities (Fig. 4e-h). Interestingly, high Ca concentrations in the biomass also increased litter quality (path coefficient of 0.32). In addition, N enrichment and plant species richness had opposite (positive and negative, respectively) indirect effects on litter quality because they had opposing effects on the N content of the biomass.

Plant species richness increased soil quality, as observed in the mixed models (see Fig. 1b). However, this effect was not direct or through effects on soil fauna, but was indirect and mediated by a change in microclimatic conditions: increased plant cover in diverse communities presumably increased soil moisture which increased the decomposition rate. N enrichment also increased soil quality indirectly, through a change in the microclimatic conditions.

Plant functional composition also altered soil quality through changes in microclimatic conditions. Communities with high Ca contents had higher plant cover and therefore higher soil quality. Ca-rich communities were dominated by herbs, which would explain this increase in cover, as herbs established better than grasses at the start of the experiment probably due to higher drought resistance. Surprisingly, however, biomass N was negatively related to plant cover. This can be explained either by a larger investment of the more productive plants in structural tissues (higher fibre contents and a dilution of biomass N content), or by the dry conditions in the first year of the experiment, which allowed the conservative species (with low N contents) to establish better than faster growing species (see Figure S8). Ca also had a direct negative effect on soil quality. Ca therefore had opposing effects on total decomposition through its effects on litter quality (positive) and effects on soil quality (negative), with a total positive effect of 0.20. Biomass N increased, and LDMC and SLA decreased, the effect of the macrofauna on decomposition, i.e. the relative differences in decomposition rate in big compared to small bags (coeff. 0.31; −0.22 and −014 respectively). However, the change in the effect of the soil fauna did not influence soil quality (i.e. there is no path between soil fauna and soil quality).

Plant functional diversity had no significant effect on decomposition, despite the increase in the soil quality effect in mixed communities plots found in the linear models (Fig. 1b). According to the SEM, this effect seems to be mediated by mass-ratio (community-weighted traits) rather than functional diversity effects per se. Functional diversity increased with species richness, which can also be due to the coding of monoculture as zero diversity.

## Discussion

Here we disentangled the key drivers of litter decomposition by using data from several litter bag experiments to compare the effects of soil and litter quality on decomposition. We use a new approach to combine data from three types of litter bag in an experiment manipulating the direct (increase in soil N) and indirect (diversity and functional composition change) effects of N enrichment. Our results show that both litter and soil quality affect overall decomposition, but that litter quality is most important. The key determinant of litter quality was the functional composition of the plant community, which played a bigger role than plant species richness or functional diversity. It was important to consider effects of multiple mechanisms and pathways because we found that some factors had contrasting effects on soil and litter quality (like Ca), or contrasting direct and indirect effects (species richness, biomass N), meaning that we would have missed many effects if we had looked only at their overall effects on decomposition. Therefore, N enrichment increases decomposition, mostly through indirect effects arising from a shift in functional composition towards faster growing plant species which produce easily decomposable litter.

### The relative importance of litter and soil quality in determining decomposition

The overall decomposition rate was more influenced by litter quality than by soil quality in our experiment (see Figure 3). This result agrees with studies in multiple biomes showing that litter traits are more important than the complexity of the decomposer community (García-Palacios, Maestre, Kattge, & Wall, 2013) or soil properties in determining decomposition (García-Palacios et al., 2016b). However, other studies in boreal forests experiencing long term N enrichment have found opposing patterns (Maaroufi et al., 2017). Part of this variation between the outcomes of these studies might be explained by differences in the relative importance of litter versus soil quality across biomes. We might expect that soil quality would be more important in unproductive ecosystems, where soil biota are expected to react more strongly to a change in microclimatic conditions (Blankinship et al., 2011). The soil quality effect could also be stronger when N enrichment leads to a decrease in soil pH, which reduces soil community diversity and abundance (Chen, Lan, Hu, & Bai, 2015; Tian & Niu, 2015). These previous studies also used different approaches to quantify litter and soil effects on decomposition and some of the variation among them may arise because they analysed different litter traits or incorporated different measures of the soil community. By combining our different litter bag experiments, we integrate all aspects of litter quality and soil quality together, allowing us to robustly test for their relative importance without the need for a complete list of all the litter and soil properties that could affect decomposition. Further studies using our approach could compare the effects of soil and litter quality on decomposition across environmental gradients to determine the global importance of these factors in determining litter decomposition.

### Functional composition is the main driver of litter quality

The main determinants of litter quality in our experiment were related to the leaf economics spectrum. Plant communities with an N-rich biomass, low fibre content and low LDMC produced the most degradable material because this type of litter is easier for the soil fauna to break down. This result agrees with a large body of literature showing that litter quality relates to leaf traits indicating a fast growth strategy, like high SLA and biomass N, low LDMC, as well as low fibre content (Cornwell et al., 2008; Reich, 2014; Freschet, Aerts, & Cornelissen, 2012). Interestingly, in our experiment, we found that nutrient contents (N and Ca) were about twice as important as structural components (LDMC and fibres) in determining litter quality (combined path coefficients of 0.61 for nutrients and - 0.36 for structure). Effects of N have been shown in many studies (Garnier et al., 2004; Cornwell et al., 2008) and as pointed out in Mládková, Mládek, Hejduk, Hejcman, and Pakeman (2018), Ca and Mg content (which were highly correlated in our case) also indicate a better digestibility and a higher decomposability of the litter (García-Palacios et al., 2016a). Ca and Mg are key components of invertebrate diets and can therefore increase their abundance (National Research Council, 2005), which may explain their positive effects on decomposition. However, a high Ca content did not increase the effect of macrofauna on decomposition perhaps suggesting that high Ca is also important for microbes. In addition to the nutrients, litter structural components were important in determining decomposition. We found that fibre content was important alongside LDMC in determining decomposition which suggests that there are several aspects of plant structure that matter. The fibre content, measured in bulk biomass, added complementary information on structure, as some species had a low LDMC but still produced fibrous stems (see Figure S9). We did not measure plant defence compounds such as tannins and phenolics, which can also be important determinants of litter quality (Hättenschwiler & Jørgensen, 2010), however, these may correlate strongly with SLA if growth-defence trade-offs are widespread (Blumenthal et al., 2009). Overall, our results show that nutrients and structure are the key determinants of litter quality but that several different aspects are important and should be considered, as single traits may not provide adequate proxies of overall litter quality.

Litter diversity, calculated from the diversity of functional traits of the species present in the plot, did not have any effect on litter quality. Functional diversity might be of importance only in communities containing legumes, where a transfer of nutrients from the N-rich legume litter to more recalcitrant litter can increase decomposition (Handa et al., 2014). Our experimental design, which did not include legumes, may therefore have underestimated the effects of diversity on decomposition rates. Our results do however, agree with other studies using tree leaf litter which showed that functional composition is usually a good predictor of litter decomposition rate and that functional diversity is of secondary importance (see Finerty et al., 2016 and Bílá et al., 2014).

### Soil quality and soil fauna effects are indirectly mediated by biomass Ca content and microclimate

Soil quality also affected the overall decomposition rate, although it was less important than litter quality. Soil quality was influenced by two factors: biomass Ca and microclimatic conditions. We observed no direct effect of N enrichment, plant species richness, functional diversity or soil fauna on soil quality, all their effects were mediated through changes in plant cover (microclimate; see Figure 2). The key indicator of increased plant cover was biomass N, which suggests that a decrease in plant cover under N enrichment could decrease soil decomposition potential by decreasing humidity. N addition had both direct (positive) and indirect effects (through increasing the negative effect of biomass N) on plant cover. As microclimate had no impact on the relative effect of macro vs. microfauna it seems likely that an increase in humidity was of equal importance for all soil decomposers. In contrast to the positive effects of microclimate, biomass Ca reduced soil quality. This means that plant communities producing more digestible litter, with a higher Ca (and/or Mg) content, were growing on a soil which was poor at decomposing standard litter. Since we used a fairly recalcitrant standard litter, this result could indicate that inputs of Ca-rich litter stimulated soil communities that were less effective at decomposing recalcitrant litter. Enzymes responsible for the breakdown of resistant material have been shown to be inhibited under N enrichment (Carreiro, Sinsabaugh, Repert, and Parkhurst (2000), but see Sinsabaugh (2010)). Our results may indicate that these enzymes are also inhibited by inputs of Ca-rich litter. Our use of one standard material may therefore have underestimated some effects if there are strong interactions between litter and soil quality and future studies could consider using a range of standard litters. The various direct and indirect effects of N enrichment therefore had opposing effects on soil quality: a loss of species diversity, expected under N enrichment, would reduce soil quality but this effect would be compensated for by a direct increase of plant cover under fertilisation.

The relative effect of macrofauna on decomposition increased with biomass N and decreased with LDMC. The macro and mesofauna contribution to decomposition was higher, relative to the effect of microfauna, when litter contained more easily degradable material. This means that high litter quality either increased the abundance of macrofauna, such as earthworms and isopods, or their efficiency in breaking down litter. Little is known about how a change in litter quality alters the effect of different soil fauna on decomposition but we can hypothesise that macrofauna are more active when feeding on higher quality litter because they actively forage for nutrients and make them available for microorganisms (see Smith & Bradford, 2003).

Our study used a new experimental and analytical approach to disentangle the complex drivers of litter decomposition. However, some issues need to be considered and the most important of these is probably the relatively early stage of the experiment. Overall, the lower importance of soil quality compared to litter quality for decomposition indicates either that litter quality is indeed more important than soil quality, or that the effects of N enrichment, diversity and functional composition take longer to fully change soil communities (Eisenhauer et al., 2011; Boeddinghaus et al., 2019). In particular, we might expect the plant species richness effect on decomposition to become more important in longer experiments, as the soil biotic community becomes more closely linked to the aboveground community (Eisenhauer, Reich, & Scheu, 2012). The drivers of decomposition might therefore change as communities re-assemble above and belowground.

In our experiment we used green litter, as green litter decomposition is an important process in grasslands managed by mowing and very little senescent plant material is present in these grasslands. However, the factors determining decomposition of dead litter may differ. Due to its higher fibre to nutrient ratio, dead litter would have taken more time to decompose and the relative importance of litter quality compared to soil quality might have been lower. Although green litter accounts for a large part of the decomposed material in semi-natural grasslands, the decomposition of dead litter is also important and separate studies would need to explore its drivers. In addition, we measured litter mass loss after 2.5 months of decomposition. While some litter bags were almost empty at the end of the experiment, we have to keep in mind that the results represent a snapshot of the decomposition process, for some plots only the early stage of decomposition. It would be interesting to determine the drivers of litter decomposition at different stages of decomposition as the relative importance of soil and litter quality, and the factors determining them, might change over time (Smith & Bradford, 2003).

## Conclusion

Decomposition was more strongly affected by litter quality rather than soil quality under N enrichment. Aboveground plant traits related to structural composition as well as nutrient concentrations were major determinants of high litter quality. This suggests that several traits are needed to properly characterise litter quality and that stem structural composition should be considered alongside leaf traits. Soil quality was mainly affected by microclimatic conditions, driven by changes in plant cover. Our study suggests that, at least for the early stages of plant material decomposition, N enrichment will directly increase decomposition rates by increasing litter N content and by increasing biomass which promotes a microclimate favouring high soil faunal activity. It will indirectly affect decomposition through a shift in plant functional composition towards faster growing species, which will increase litter quality, and through a loss in plant species richness, which would mainly decrease soil quality through a reduction in plant cover. The relative importance of different drivers of decomposition under N enrichment might vary between ecosystems and further studies could use our approach to quantify the relative importance of soil and litter quality in different contexts. Nevertheless, the large effect of plant functional composition, seen in both biomass nutrients and structural components, indicates that it is among the major drivers to take into consideration when assessing overall N enrichment effects on decomposition.

## Acknowledgments

We are grateful to Jon Lefcheck and Jim Grace for the very helpful exchange of emails about structural equation models. Pablo Garcia Palacios kindly reviewed an earlier version of the manuscript. We would like to thank the technicians and the team of 100 helpers of the PaNDiv Experiment, as well as the students helping in Münster. This study was supported by funding of the Swiss National Science Foundation. SS was supported by the Spanish Government under a Ramón y Cajal contract (RYC-2016-20604).

## Authors’ contribution

NP, SC and EA designed and set up the PaNDiv Experiment. NP and SC collected the data. NP, NH, VHK and TK processed and analysed the NIRS samples. NP analysed the data and wrote the first manuscript with the substantial input from EA, SS and SC. All authors contributed to revisions of the manuscript.

## Data accessibility

Once this manuscript is accepted, all the relevant data will be archived in figshare (https://figshare.com/).

