## Supporting information for "Decomposition disentangled: a test of the multiple mechanisms by which nitrogen enrichment alters litter decomposition"

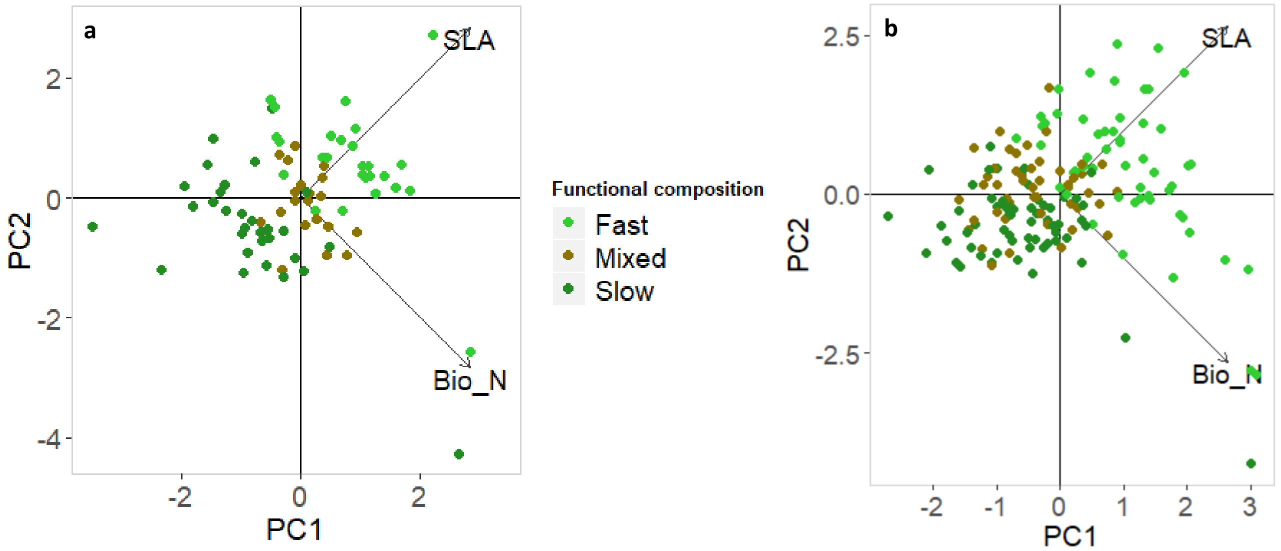


**Figure S1:** Community traits Specific Leaf Area (SLA) and biomass nitrogen for our 168 communities, divided by fast, mixed and slow plots. **a**) The sown community traits calculated using control monocultures SLA and biomass N (expected values). **b**) The community weighted mean traits calculated using control monocultures and species percentage cover per plot (realised values).

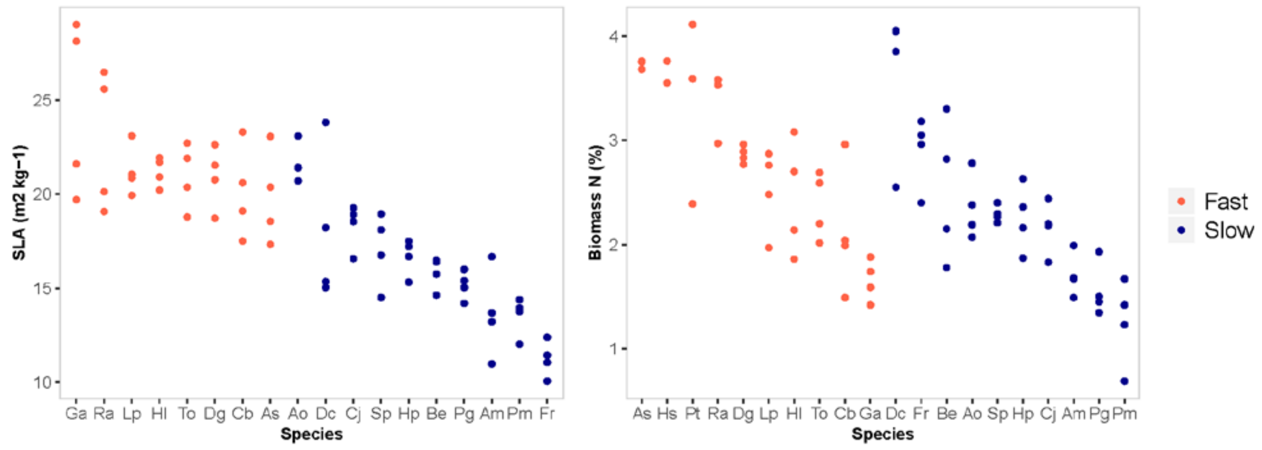


**Figure S2:** Specific leaf area and biomass N content measured per plot in the four monocultures. List of species:

Fast growing:

*Anthriscus sylvestris* (As), *Crepis biennis* (Cb), *Dactylis glomerata* (Dg), *Galium album* (Ga), *Holcus lanatus* (Hl), *Heracleum sphondylium* (Hs), *Lolium perenne* (Lp), *Poa trivialis* (Pt), *Rumex acetosa* (Ra) and *Taraxacum officinale* (To).

Slow growing:
*Achillea millefolium* (Am), *Anthoxanthum odoratum* (Ao), *Bromus erectus* (Be), *Centaurea jacea* (Cj), *Daucus carotta* (Dc), *Festuca rubra* (Fr), *Helictotrichon pubescens* (Hp), *Prunella grandiflora* (Pg), *Plantago media* (Pm) and *Salvia pratensis* (Sp).

**Table S1**: The PaNDiv Experiment manipulates species diversity, functional composition, nitrogen enrichment and foliar fungal pathogens in a full factorial design. The 4 and 8 species levels consist per treatment in 10 fast, 10 mixed and 10 slow communities. We were growing 10 fast and 10 slow monocultures and 4 replicates of the 20 species together. These 84 communities were sown in a control plot, with nitrogen, with fungicide, and with both nitrogen and fungicide. The total number of plots summed up to 336 divided in four blocks. Each block contained all 84 combination with a randomly attributed treatment.

| **Manipulation** | **Levels** | |
|---|---|---|
| Species diversity (SD) | Monoculture | *20 combinations* |
| | 4 species | *30 combinations* |
| | 8 species | *30 combinations* |
| | 20 species | *4 replicates* |
| Functional composition (FC) | Fast growing | |
| | Slow growing | |
| | Mix of both | |
| Nitrogen enrichment (Ni) | 100 kg N $ha^{-1}$ $y^{-1}$ | |
| | no fertilisation | |
| Fungicide (Fz) | fungicide | |
| | no fungicide | |

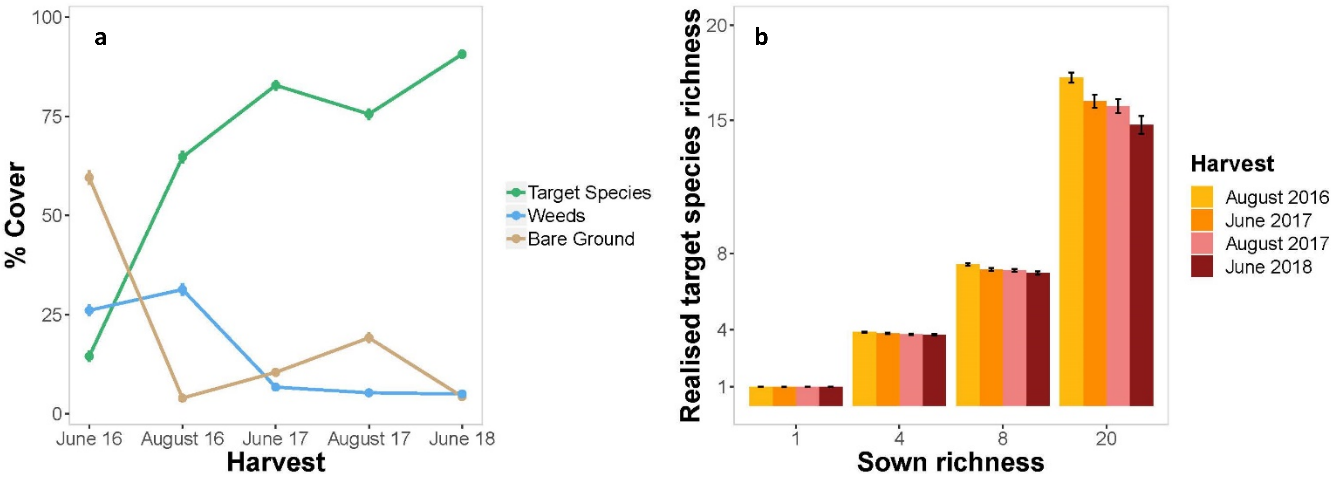


**Figure S3:** Field establishment across time. Percentage cover of target species, weeds and bare ground (**a**). Number of target species within each diversity level over time (**b**).

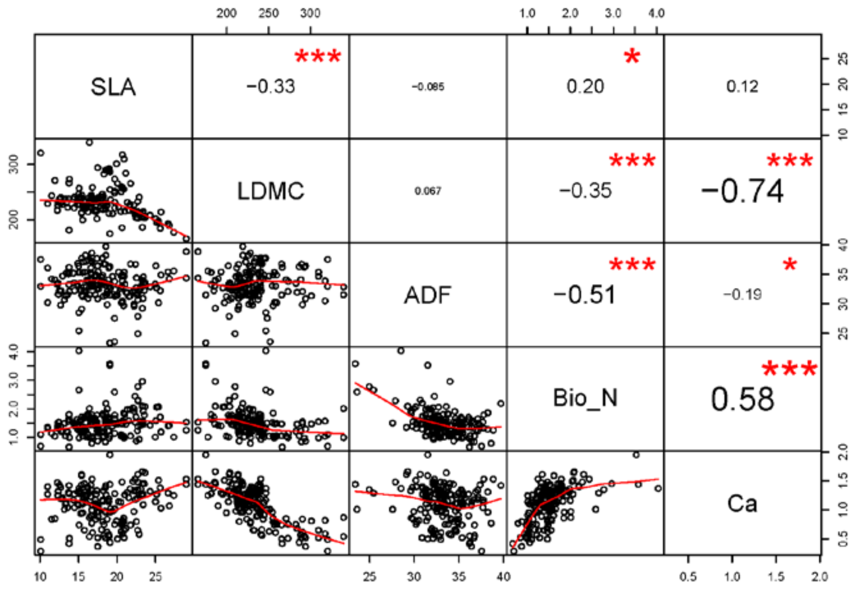


**Figure S4:** Spearman correlations between the community weighted mean measured traits specific leaf area and leaf dry matter content, and the NIRS (near infrared reflectance spectroscopy) analysed elements in the biomass, using the R package "PerformanceAnalytics". ADF = Acid Detergent Fibres. Calcium and LDMC do have a high correlation (-0.74) and we have to interpret results including both variables with caution, as we may underestimate their effects.

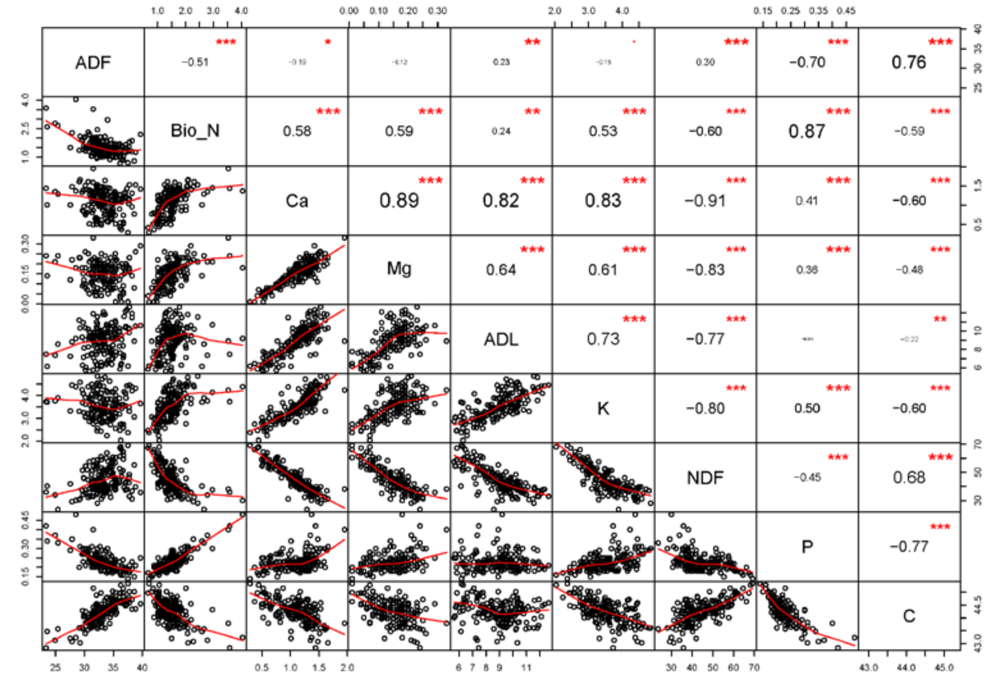


**Figure S5:** Spearman correlations between the NIRS analysed elements in the biomass, using the R package "PerformanceAnalytics". ADF = Acid Detergent Fibres; ADL = Acid Detergent Lignin; NDF = Neutral Detergent Fibres. We used ADF, biomass N and Ca in the structural equation model.

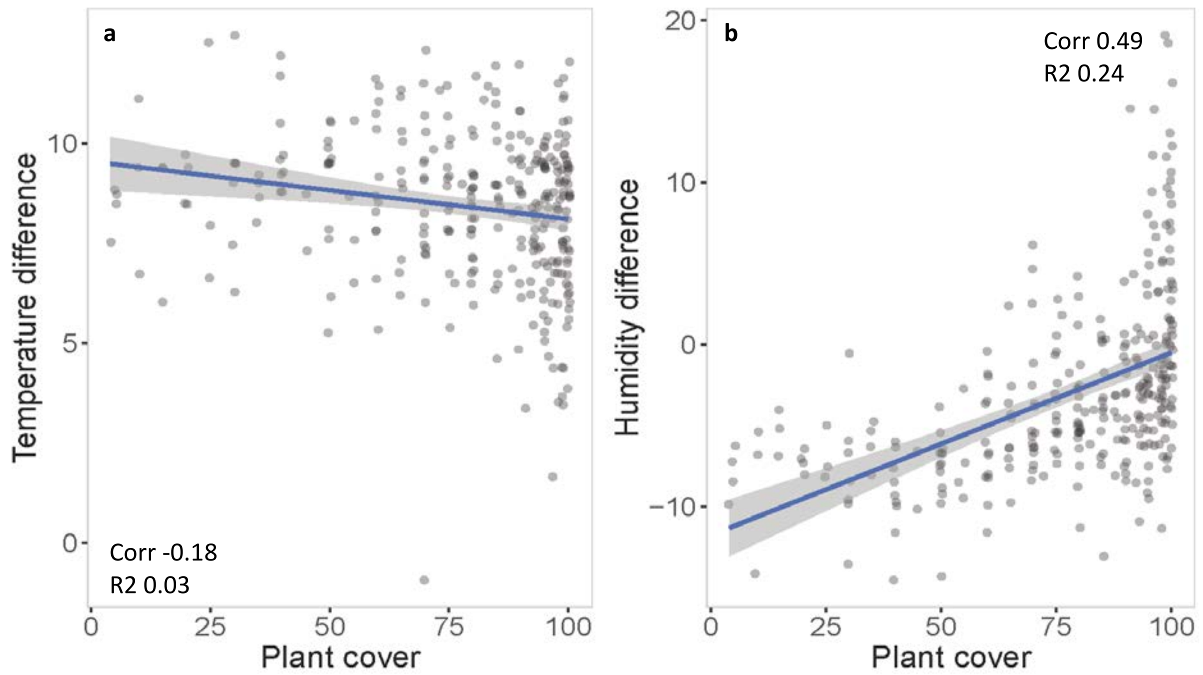


**Figure S6:** Effect of plant cover on microclimatic conditions: **a)** temperature and **b)** humidity. Plant cover was estimated in August 2018. The temperature and humidity were measured in July and August 2018 on each plot for minimum 2 days over a period of four weeks. Each point represents the mean deviation of the microclimatic temperature in the vegetation from the values recorded at the closest meteorological Station in Zollikhofen (ca. 4km away).

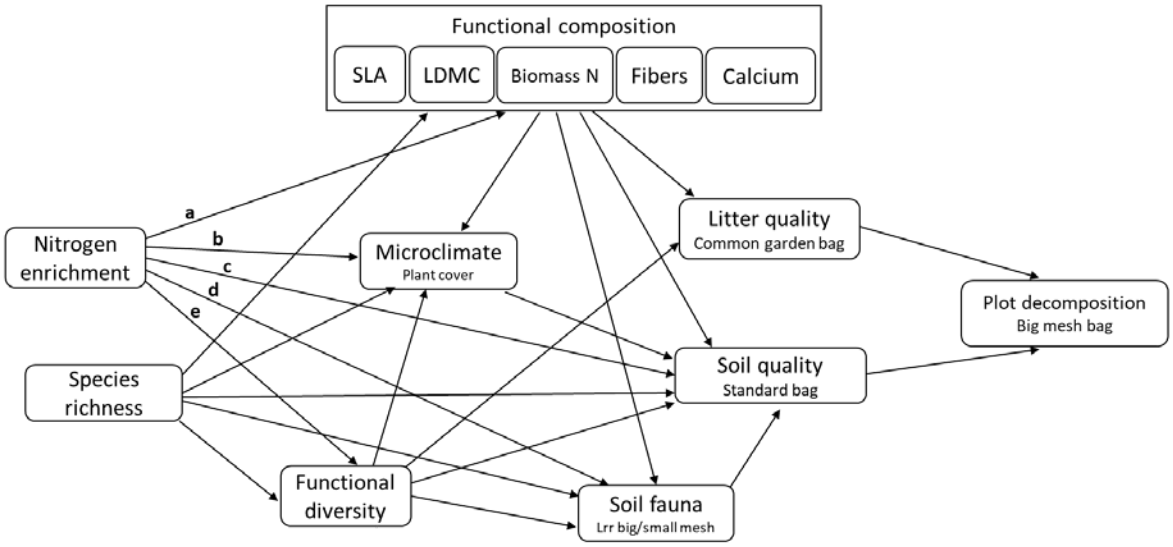


**Figure S7:** A priori model structure for the Structural Equation Model analysis. Hypothesis:

N enrichment shifts functional composition (**a**) and decreases functional diversity (**b**), affecting litter mediated decomposition. N enrichment affects soil quality through an increase of microclimatic conditions (**c**), shifts in soil fauna relative effect (**d**), or any other effect not taken into account here (**e**) for instance a change in available nitrogen. We expect species richness to influence the same variables. We included all these arrows because we know little about the importance of the different processes.

Functional composition and functional diversity affect litter mediated decomposition by changing litter decomposability, and soil through effects on microclimate, soil fauna or any other direct effect materialised by a direct arrow, as for N and SR.

The plot decomposition rate is affected via both litter mediated and soil mediated effect.

The final model included some correlations to increase the model fit, and because they are related to potential unmeasured variables that affected the outcome of the multiple experiments plugged in the model. We included a correlation between biomass nitrogen and functional diversity and between LDMC and functional diversity because of the way we coded monocultures (see main text). We also included a correlation between plot decomposition and soil fauna, because the fauna variable is calculated from plot decomposition. The last correlation is between litter mediated and plot decomposition. In these two experiments, we used the same litter material (see main text).

**Table S2:** We fitted the model using piecewiseSEM (Lefcheck, 2016) another package for Structural Equation Model computing several linear mixed effect models together. Unlike the lavaan package, it enabled us to include "Combination" as a random term. Each plot contains a different species combination, grown as a control or with nitrogen addition. If the random term is not included, it increases the species richness effect. Comparing lavaan and piecewiseSEM outputs informed us about the importance of this potential bias. We decided to use lavaan in the end because the differences between the packages were not major and because of lavaan's wide use. At the same time, the recent update of the piecewiseSEM package with unfixed issues made it difficult to work with. The results presented here are calculated using the earlier version 1.2.1.

When using the same initial model as for the lavaan code, the piecewiseSEM model was rejected (fit using fischer.c, p.value significant). The model fit improved when we added three residual covariances (p.value 0.117): between plot decomposition and calcium, biomass N, and fibres. However, the path coefficients did not change at all when we added these covariances.

In the piecewiseSEM output, three paths became non-significant: Litter quality ~ LDMC, Soil fauna ~ SLA and Microclimate ~ Species richness. Two other became marginally significant: biomass N ~ Species richness, Litter quality ~ Fibres. The differences are highlighted in the table hereafter. The estimates of soil and litter quality on plot decomposition were lower in the piecewiseSEM output but their relative importance stays the same. On the other hand, the effect of species richness on biomass N and on Functional diversity was underestimated in lavaan. Although the model output differed slightly between the two packages, the overall conclusions of our study remains the same.

| Response | Predictor | Estimate | Std.error | p.value | |
|---|---|---|---|---|---|
| Plot decomposition | Litter quality | 0.652 | 0.061 | 0.000 | *** |
| | Soil quality | 0.130 | 0.060 | 0.033 | * |
| Litter quality | Biomass N | 0.335 | 0.085 | 0.000 | *** |
| | Ca | 0.323 | 0.100 | 0.002 | ** |
| | Fibres | -0.120 | 0.062 | 0.060 | |
| | LDMC | -0.138 | 0.101 | 0.180 | |
| | SLA | -0.056 | 0.070 | 0.430 | |
| | Functional diversity | -0.012 | 0.068 | 0.863 | |
| Soil quality | Microclimate | 0.296 | 0.099 | 0.004 | ** |
| | Ca | -0.390 | 0.148 | 0.011 | * |
| | LDMC | -0.204 | 0.133 | 0.132 | |
| | Fibres | -0.065 | 0.090 | 0.475 | |
| | Soil fauna | 0.041 | 0.080 | 0.610 | |
| | Species richness | 0.047 | 0.088 | 0.639 | |
| | SLA | -0.035 | 0.085 | 0.681 | |
| | N | -0.033 | 0.082 | 0.685 | |
| | Biomass N | -0.053 | 0.141 | 0.714 | |
| | Functional diversity | -0.014 | 0.094 | 0.885 | |
| Soil fauna | LDMC | -0.307 | 0.133 | 0.024 | * |
| | Biomass N | 0.277 | 0.123 | 0.027 | * |
| | SLA | -0.111 | 0.085 | 0.198 | |
| | N | -0.067 | 0.079 | 0.402 | |

| | | | | | |
|---|---|---|---|---|---|
| | Ca | -0.111 | 0.144 | 0.447 | |
| | Species richness | 0.046 | 0.089 | 0.647 | |
| | Functional diversity | -0.030 | 0.095 | 0.760 | |
| | Fibres | 0.002 | 0.091 | 0.981 | |
| SLA | N | 0.030 | 0.024 | 0.207 | |
| | Species richness | 0.039 | 0.148 | 0.791 | |
| Biomass N | N | 0.138 | 0.058 | 0.020 | * |
| | Species richness | -0.226 | 0.120 | 0.066 | |
| Fibres | Species richness | 0.098 | 0.100 | 0.347 | |
| | N | 0.054 | 0.073 | 0.460 | |
| Ca | N | -0.029 | 0.062 | 0.645 | |
| | Species richness | 0.045 | 0.120 | 0.708 | |
| LDMC | Species richness | -0.232 | 0.140 | 0.101 | |
| | N | -0.030 | 0.039 | 0.442 | |
| Functional diversity | Species richness | 0.837 | 0.117 | 0.000 | *** |
| | N | -0.015 | 0.021 | 0.466 | |
| Microclimate | Biomass N | -0.710 | 0.098 | 0.000 | *** |
| | N | 0.264 | 0.063 | 0.000 | *** |
| | Ca | 0.447 | 0.115 | 0.000 | *** |
| | SLA | -0.120 | 0.068 | 0.083 | |
| | Species richness | 0.148 | 0.071 | 0.106 | |
| | Fibres | -0.110 | 0.073 | 0.141 | |
| | LDMC | -0.128 | 0.106 | 0.236 | |
| | Functional diversity | 0.091 | 0.076 | 0.245 | |

| Covariances | | Estimate | p.value | |
|---|---|---|---|---|
| Biomass N | Ca | 0.631 | 0.000 | *** |
| | Functional diversity | 0.025 | 0.378 | |
| | LDMC | -0.403 | 1.000 | |
| | Fibres | -0.466 | 1.000 | |
| Fibres | Ca | -0.162 | 0.980 | |
| LDMC | Fibres | 0.098 | 0.109 | |
| | Functional diversity | -0.057 | 0.764 | |
| | Ca | -0.700 | 1.000 | |
| SLA | Ca | 0.231 | 0.002 | ** |
| | Biomass N | 0.177 | 0.012 | * |
| | Fibres | -0.059 | 0.772 | |
| | LDMC | -0.636 | 1.000 | |
| Plot decomposition | Soil fauna | 0.643 | 0.000 | *** |
| | Litter quality | 0.610 | 0.000 | *** |
| | Biomass N | 0.249 | 0.001 | *** |
| | Ca | 0.248 | 0.001 | *** |
| | Fibres | -0.216 | 0.997 | |

**Table S3**: Linear mixed effect models standardised output. p-values were derived by dropping a term from the model and comparing models with and without the term of interest. Model simplification was done step wise and main effects that are part of significant interactions were therefore not dropped from the model, they are indicated as "marginal".

| Simple model | Factor | Estimate | Std.Error | Pr(Chi) |
|---|---|---|---|---|
| Small Bags | (Intercept) | -0.04 | 0.11 | |
| | Nitrogen | 0.11 | 0.03 | 0.002 |
| | Litter august | 0.27 | 0.09 | 0.006 |
| Medium Bags | (Intercept) | -0.06 | 0.07 | |
| | Nitrogen | 0.13 | 0.04 | 0.001 |
| Big Bags | (Intercept) | 0.76 | 0.19 | |
| | Nitrogen | 0.16 | 0.05 | 0.004 |
| | Litter august | 0.66 | 0.17 | <0.001 |
| Common Garden | (Intercept) | 1.57 | 0.16 | |
| | Nitrogen | 0.12 | 0.05 | 0.02 |
| | Litter august | 0.88 | 0.18 | <0.001 |
| Standard Litter | (Intercept) | -1.15 | 0.09 | |
| | Species richness | 0.11 | 0.04 | 0.015 |

| Combined model | Factor | Estimate | Std.Error | |
|---|---|---|---|---|
| Bag Type | (Intercept) | 0.77 | 0.15 | |
| | Nitrogen | 0.11 | 0.03 | 0.002 |
| | Mixed Plots | -0.11 | 0.16 | marginal |
| | Slow Plots | -0.26 | 0.15 | |
| | Litter quality | 0.65 | 0.12 | |
| | Soil quality | -1.78 | 0.12 | |
| | Litter august | 0.50 | 0.11 | <0.001 |
| | Mixed Plots * Litter quality | 0.02 | 0.18 | 0.035 |
| | Slow Plots * Litter quality | 0.12 | 0.17 | |
| | Mixed Plots * Soil quality | 0.42 | 0.18 | |
| | Slow Plots * Soil quality | 0.47 | 0.17 | |
| Mesh size | (Intercept) | 0.57 | 0.12 | |
| | Nitrogen | 0.14 | 0.03 | <0.001 |
| | Medium bag | -0.40 | 0.05 | <0.001 |
| | Small bag | -0.54 | 0.05 | |
| | Litter august | 0.38 | 0.10 | <0.001 |

| Simple traits model | Factor | Estimate | Std.Error | |
|---|---|---|---|---|
| Small Bags | (Intercept) | -0.21 | 0.08 | |
| | Nitrogen | 0.09 | 0.03 | 0.007 |
| | LDMC | -0.12 | 0.03 | <0.001 |
| Medium Bags | (Intercept) | -0.08 | 0.06 | |
| | Nitrogen | 0.13 | 0.04 | 0.001 |
| | Species richness | 0.01 | 0.06 | marginal |
| | LDMC | -0.21 | 0.06 | |

| | | | | |
|---|---|---|---|---|
| | Species richness * LDMC | -0.15 | 0.07 | 0.033 |
| Big Bags | (Intercept) | 0.76 | 0.16 | |
| | Nitrogen | 0.13 | 0.05 | 0.019 |
| | LDMC | -0.35 | 0.06 | <0.001 |
| | Litter august | 0.65 | 0.18 | <0.001 |
| Common Garden | (Intercept) | 1.45 | 0.13 | |
| | Nitrogen | 0.11 | 0.05 | 0.040 |
| | LDMC | -0.47 | 0.06 | <0.001 |
| | Functional diversity | -0.16 | 0.07 | 0.017 |
| | Litter august | 0.61 | 0.18 | 0.001 |
| | Initial litter | -0.15 | 0.06 | 0.015 |
| Standard Litter | (Intercept) | -1.15 | 0.10 | |
| | Functional diversity | 0.10 | 0.04 | 0.007 |

| Combined traits model | Factor | Estimate | Std.Error | |
|---|---|---|---|---|
| Bag Type | (Intercept) | 0.61 | 0.12 | |
| | Nitrogen | 0.09 | 0.03 | 0.009 |
| | Litter quality | 0.69 | 0.06 | marginal |
| | Soil quality | -1.49 | 0.06 | |
| | LDMC | -0.39 | 0.05 | |
| | Functional diversity | -0.06 | 0.06 | |
| | Litter august | 0.42 | 0.11 | <0.001 |
| | Litter quality * LDMC | -0.05 | 0.07 | <0.001 |
| | Soil quality * LDMC | 0.49 | 0.07 | |
| | Litter quality * Functional diversity | -0.12 | 0.07 | <0.001 |
| | Soil quality * Functional diversity | 0.18 | 0.07 | |
| Mesh size | (Intercept) | 0.54 | 0.11 | |
| | Nitrogen | 0.13 | 0.03 | <0.001 |
| | Medium bag | -0.39 | 0.05 | marginal |
| | Small bag | -0.54 | 0.05 | |
| | LDMC | -0.38 | 0.05 | |
| | Litter august | 0.32 | 0.11 | 0.004 |
| | Medium bag * LDMC | 0.31 | 0.05 | <0.001 |
| | Small bag * LDMC | 0.28 | 0.05 | |

**Table S4**: Structural Equation Model standardised output using lavaan.

| Response | Predictor | Estimate | Std.Err | P(>\|z\|) |
|---|---|---|---|---|
| Plot decomposition | Litter quality | 0.933 | 0.092 | 0 |
| | Soil quality | 0.192 | 0.042 | 0 |
| Litter quality | Functional diversity | -0.019 | 0.05 | 0.705 |
| | SLA | -0.023 | 0.05 | 0.645 |
| | Biomass N | 0.344 | 0.077 | 0 |
| | Fibres | -0.163 | 0.055 | 0.003 |
| | Ca | 0.262 | 0.084 | 0.002 |
| | LDMC | -0.197 | 0.08 | 0.014 |
| Soil quality | Soil fauna | 0.045 | 0.083 | 0.59 |
| | N enrichment | -0.033 | 0.079 | 0.674 |
| | Microclimate | 0.328 | 0.106 | 0.002 |
| | SLA | -0.035 | 0.082 | 0.669 |
| | Biomass N | -0.057 | 0.149 | 0.701 |
| | Fibres | -0.066 | 0.089 | 0.453 |
| | Ca | -0.389 | 0.142 | 0.006 |
| | LDMC | -0.206 | 0.13 | 0.112 |
| | Functional diversity | -0.014 | 0.092 | 0.877 |
| | Species richness | 0.047 | 0.085 | 0.578 |
| Soil fauna | Species richness | 0.038 | 0.059 | 0.513 |
| | N enrichment | -0.101 | 0.052 | 0.054 |
| | SLA | -0.146 | 0.063 | 0.021 |
| | Biomass N | 0.331 | 0.1 | 0.001 |
| | Fibres | 0.063 | 0.07 | 0.369 |
| | Ca | -0.088 | 0.108 | 0.414 |
| | LDMC | -0.269 | 0.101 | 0.008 |
| | Functional diversity | -0.031 | 0.07 | 0.659 |
| SLA | N enrichment | 0.021 | 0.079 | 0.787 |
| | Species richness | -0.008 | 0.079 | 0.922 |
| Biomass N | N enrichment | 0.14 | 0.07 | 0.047 |
| | Species richness | -0.141 | 0.071 | 0.046 |
| Fibres | N enrichment | 0.052 | 0.077 | 0.5 |
| | Species richness | 0.07 | 0.078 | 0.364 |
| Ca | N enrichment | -0.02 | 0.079 | 0.799 |
| | Species richness | 0.031 | 0.079 | 0.693 |
| LDMC | N enrichment | -0.042 | 0.078 | 0.589 |
| | Species richness | -0.128 | 0.078 | 0.1 |
| Functional diversity | Species richness | 0.507 | 0.066 | 0 |
| | N enrichment | -0.022 | 0.066 | 0.745 |
| Microclimate | Species richness | 0.134 | 0.062 | 0.031 |
| | N enrichment | 0.239 | 0.056 | 0 |

| | | | |
|---|---|---|---|
| SLA | -0.108 | 0.059 | 0.07 |
| Biomass N | -0.696 | 0.094 | 0 |
| Fibres | -0.1 | 0.065 | 0.123 |
| Ca | 0.403 | 0.101 | 0 |
| LDMC | -0.117 | 0.095 | 0.216 |
| Functional diversity | 0.084 | 0.068 | 0.215 |

| Covariances | | Estimate | Std.Err | P(>\|z\|) |
|---|---|---|---|---|
| Biomass N | LDMC | -0.385 | 0.075 | 0 |
| | Ca | 0.548 | 0.082 | 0 |
| | Fibres | -0.455 | 0.077 | 0 |
| Fibres | Ca | -0.179 | 0.078 | 0.023 |
| | LDMC | 0.11 | 0.075 | 0.145 |
| Ca | LDMC | -0.758 | 0.097 | 0 |
| SLA | LDMC | -0.325 | 0.081 | 0 |
| | Biomass N | 0.232 | 0.072 | 0.001 |
| | Fibres | -0.093 | 0.078 | 0.234 |
| | Ca | 0.131 | 0.08 | 0.1 |
| Plot decomposition | Soil fauna | 0.394 | 0.056 | 0 |
| | Litter quality | -0.331 | 0.063 | 0 |
| LDMC | Functional diversity | -0.155 | 0.041 | 0 |
| Biomass N | Functional diversity | -0.109 | 0.039 | 0.006 |

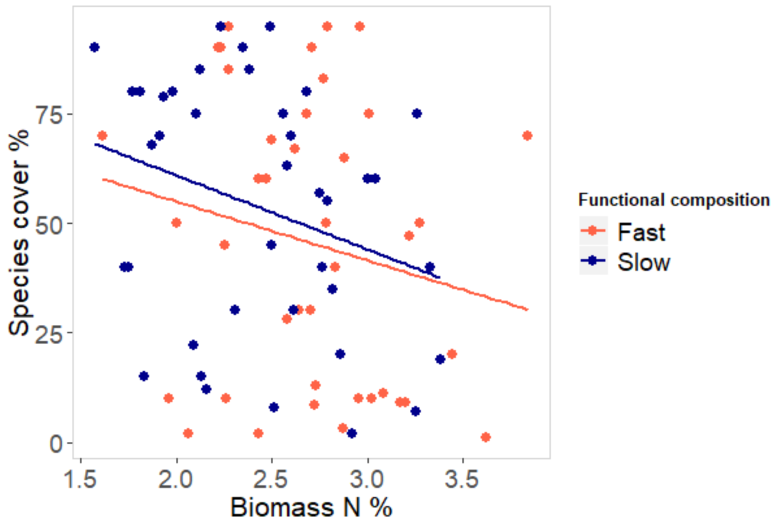


**Figure S8**: Correlation in August 2016 between species percentage cover and biomass N content (%) in the monocultures fast and slow growing plots. At the beginning of the experiment, species with low biomass N content established better than species with high biomass N content.

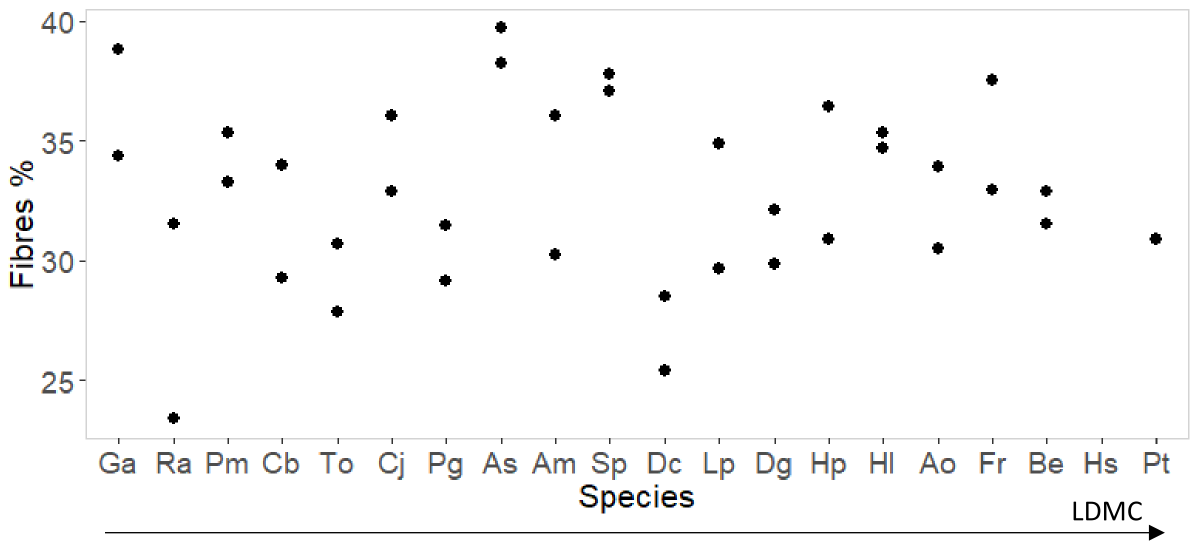


**Figure S9**: Percentage fibre content (acid detergent lignin) in the monoculture plots biomass. Species are ranked from left to right in increasing order of LDMC. Some species like *Galium album* or *Plantago media* present both a relatively low LDMC and a large fibre content.
